# Hepatitis C Virus Remodels Lipid Droplets to Promote Mitochondrial Fatty Acid Accumulation and Metabolic Activation

**DOI:** 10.64898/2026.07.09.737644

**Authors:** Imaan Muhammad, Kaci Craft, Shaokai Pei, Kaia Conte, Junyan Li, Shaolei Teng, Ruth Cruz-cosme, Shaomin Yang, Yanjin Zhang, Qiyi Tang

**Affiliations:** Department of Microbiology, Howard University College of Medicine, Washington, DC 20059; Department of Biology, Howard University, Washington, DC 20059; Guangdong Key Laboratory for Biomedical Measurements and Ultrasound Imaging, National-Regional Key Technology Engineering Laboratory for Medical Ultrasound, School of Biomedical Engineering, Shenzhen University Medical School, Shenzhen, 518060, China; Department of Veterinary Medicine, University of Maryland, College Park, Maryland

## Abstract

Hepatitis C virus (HCV) depends on host lipid metabolism and lipid droplets (LDs) for genome replication, assembly, and particle production, yet how LD structure and lipid utilization change over the course of infection remains incompletely understood. Here, we investigated the temporal remodeling of LD-associated metabolic pathways during HCV JFH-1 infection of human hepatoma Huh7 cells. HCV infection transiently increased LD fluorescence intensity at 24 hours post-infection (hpi), followed by normalization or relative loss of LD signal at later time points. Concomitantly, LDs became progressively clustered and enlargement during late infection, despite reduced protein levels of the canonical LD fusion proteins CIDEA, CIDEB, and CIDEC, suggesting that HCV-induced LD enlargement occurs through CIDE-independent mechanisms. Transcriptomic, RT-qPCR, and immunoblot analyses revealed time-dependent regulation of genes and proteins involved in LD structure, triglyceride synthesis, lipolysis, lipid uptake, and mitochondrial fatty acid utilization. Subcellular fractionation demonstrated preferential accumulation of fatty acids in mitochondrial fractions at 24–72 hpi. This redistribution was accompanied by increased oxygen consumption rate, elevated extracellular acidification, and progressive reactive oxygen species accumulation, indicating infection-associated metabolic activation and oxidative stress. Pharmacological inhibition of DGAT1-dependent LD biogenesis, LIPA-dependent lysosomal lipid hydrolysis, LIPE/HSL-dependent lipolysis, or CPT1-dependent mitochondrial fatty acid transport markedly reduced mitochondrial fatty acid accumulation and suppressed HCV-induced respiratory activity. Inhibition of LIPA or LIPE/HSL reduced both HCV RNA and core protein levels, whereas inhibition of CPT1 or DGAT1 had more pronounced effects on core protein than on viral RNA. Together, these findings support a model in which HCV dynamically remodels LDs, mobilizes LD-associated fatty acids, and redirects them toward mitochondria to support infection-associated metabolism and downstream stages of the viral life cycle. Lipid hydrolysis and mitochondrial fatty acid trafficking therefore represent potential host-directed targets for limiting HCV infection.

**SIGNIFIGANCE:** Hepatitis C virus depends on host lipid metabolism for replication, assembly, and production of infectious particles, but how it uses lipid droplets over time remains incompletely understood. This study shows that hepatitis C virus dynamically remodels lipid droplets, causing an early increase in lipid storage followed by droplet enlargement and mobilization of fatty acids during later infection. The released fatty acids preferentially accumulate in mitochondria, where they are associated with increased cellular respiration and oxidative stress. Blocking lipid droplet formation, lipid breakdown, or fatty acid transport to mitochondria reduced this metabolic response and decreased viral RNA or core protein accumulation. Inhibition of lysosomal acid lipase and hormone-sensitive lipase suppressed both viral RNA and protein levels. These findings identify lipid droplet breakdown and mitochondrial fatty acid trafficking as important host processes used by hepatitis C virus and as potential targets for host-directed antiviral intervention.

## INTRODUCTION

Hepatitis C virus (HCV) is an enveloped, positive-sense RNA virus of the *Hepacivirus* genus within the *Flaviviridae* family. HCV primarily infects hepatocytes and remains a major cause of chronic liver disease, including fibrosis, cirrhosis, and hepatocellular carcinoma ^1–3^. Although direct-acting antivirals, including glecaprevir and sofosbuvir ^4–8^, have greatly improved treatment outcomes, HCV continues to cause substantial liver-related morbidity due to ongoing transmission, delayed diagnosis, limited access to therapy for children under 3 years old, and the absence of a protective vaccine ^9–12^ . Therefore, defining the host pathways that support HCV replication and pathogenesis remains important for understanding viral persistence and identifying complementary antiviral strategies.

A defining feature of HCV infection is its close relationship with host lipid metabolism. Clinically, HCV infection is associated with dyslipidemia, altered apolipoprotein metabolism, hypobetalipoproteinemia, and hepatic steatosis ^13–15^, underscoring the broad metabolic reprogramming that accompanied infection. These alterations reflect the dependence of HCV on lipid-associated pathways throughout its life cycle. HCV circulates as lipoviroparticles containing host lipids and apolipoproteins, which facilitate interactions with lipid uptake receptors during viral entry ^16–18^. Following entry and translation at the endoplasmic reticulum (ER), viral RNA replication occurs on remodeled modified intracellular membranes, while virion assembly occurs at ER subdomains regions closely associated with lipid droplets (LDs) ^19–22^. Thus, HCV infection provides a well-established model for understanding how positive-sense RNA viruses exploit host lipid metabolism and lipid organelles to coordinate support viral replication, assembly, and particle production.

LDs are dynamic cytoplasmic organelles that store neutral lipids, triacylglycerols and cholesteryl esters, within a hydrophobic core surrounded by a phospholipid monolayer ^23–26^. LD-associated proteins, particularly perilipin family members, regulate droplet stability, lipid access, turnover, and interactions with other organelles ^27^. In hepatocytes, LDs play central roles in neutral-lipid storage, membrane biogenesis synthesis, fatty acid metabolism, and cellular energy homeostasis ^25, 28, 29^. During infection, however, viruses can remodel LD abundance, distribution of localization, composition, and turnover to establish create intracellular environments that favor replication and assembly^25, 29, 30^. LDs are particularly important for HCV infection because the viral core protein localizes to the LD surface, where it recruits viral and host factors required for contributing capsid assembly and infectious particle production^19, 30, 31^. The viral nonstructural protein NS5A is also recruited to LD-proximal ER membranes, where it coordinates the transfer of newly synthesized viral RNA to core-containing assembly sites, thereby linking genome replication to RNA packaging and virion formation^32–34^.

Although LDs are well established as platforms for HCV assembly, how HCV regulates LD abundance, turnover, and metabolic use during infection remains incompletely defined. This question is particularly important because stored lipids can be mobilized through lipolysis or lipophagy, releasing fatty acids that enter mitochondrial β-oxidation and support cellular energy production ^13, 35^. However, HCV has also been reported to impair mitochondrial fatty acid oxidation and redirect lipid metabolism toward storage and steatosis ^36, 37^. These divergent effects suggest that HCV may reshape LD metabolism in a stage- and context-dependent manner to balance lipid storage, viral assembly, and cellular energy demand. Understanding this balance may reveal host lipid pathways that are required for efficient HCV replication and pathogenesis and contribute to HCV-associated liver disease.

In this study, we investigated how HCV infection alters LD dynamics and lipid metabolic pathways in hepatocyte-derived cells. We examined whether HCV alters LD abundance over the course of infection and whether these changes are linked to viral RNA replication and particle production. We hypothesized that HCV mobilizes LD-derived lipids to generate fatty acids that support the metabolic demands of viral replication and that this process depends on coordinated regulation of LD biogenesis and lipolysis. By defining how HCV manipulates LD formation, turnover, and metabolic utilization, this study may identify host lipid pathways that can be targeted to disrupt HCV replication and limit virion production while potentially limiting HCV-associated metabolic liver injury.

## RESULTS

### 1. HCV infection transiently increases lipid droplet intensity during early infection

LDs have been closely linked to the HCV life cycle ^38^. HCV RNA replication occurs near LDs, and viral core and nonstructural proteins, including NS5A, associate with LDs. Previous studies have also reported that HCV infection induces LD accumulation ^39, 40^. To determine how HCV infection affects LD abundance over time, Huh7 cells were infected with HCV at an MOI of 1 and stained for LDs, HCV core protein, and nuclei from 24 to 96 hpi. LD fluorescence was quantified by ImageJ in more than 600 cells through 120 hpi.

At 24 hpi, HCV-infected cells exhibited abundant LD staining, with LDs distributed throughout the cytoplasm and enriched near HCV core-positive regions (Figure 1A). Quantitative analysis from more than 600 HCV-infected vs 600 mock cells confirmed that LD intensity was significantly higher in infected cells than in uninfected controls at this early time point (Figure 1B). However, this increase was no longer evident beginning at 36 hpi. From 36 to 120 hpi, LD intensity in infected cells was not significantly different from that in uninfected controls and was reduced relative to the elevated level observed at 24 hpi.

**Figure 1.**
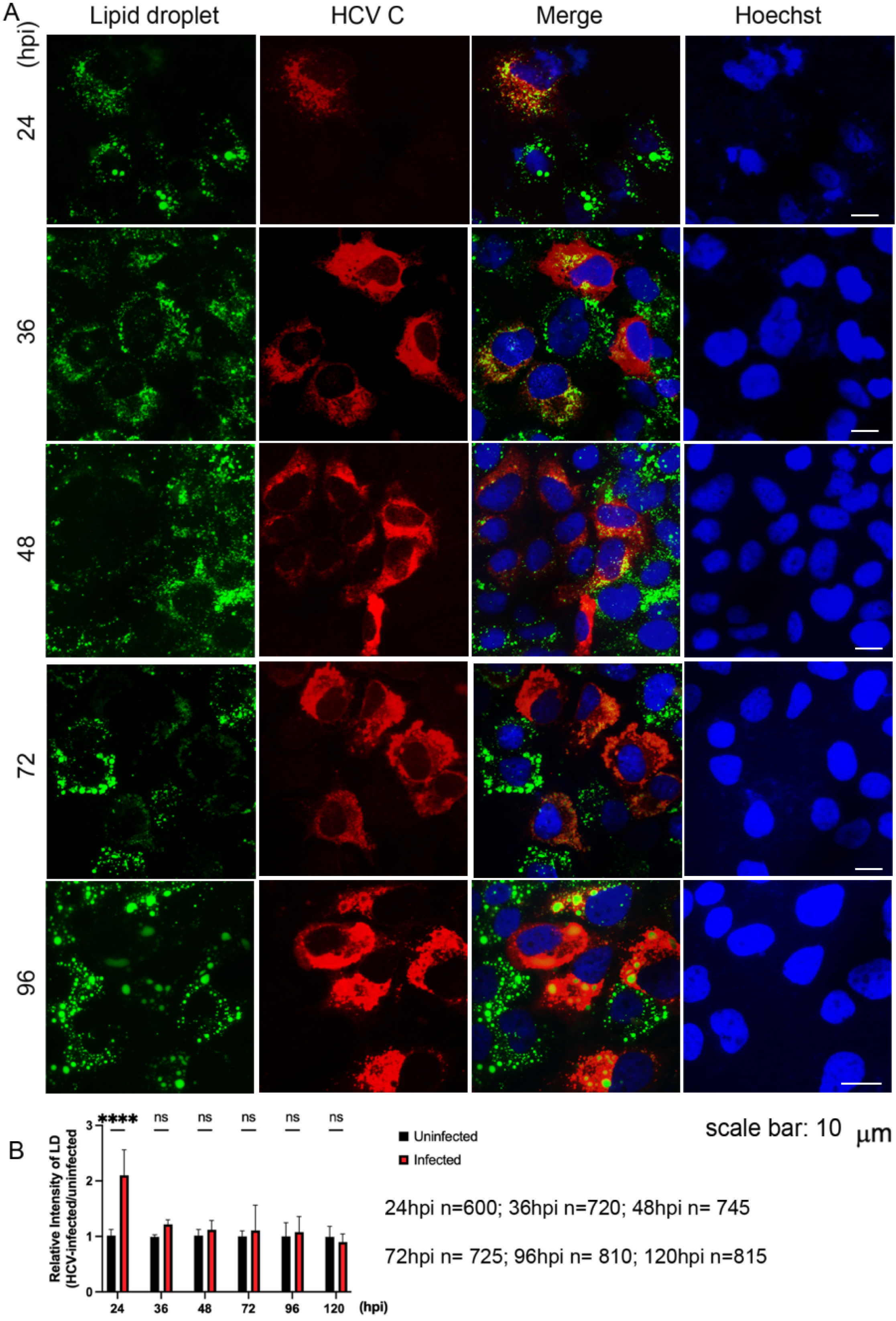
HCV-JFH1 infection transiently increases lipid droplet intensity during early infection. Huh7 cells were mock-infected or infected with HCV-JFH1 at an MOI of 1 and fixed at 24, 36, 48, 72, and 96 h post-infection (hpi). Cells were stained for lipid droplets using BODIPY 493/503 (green), HCV core protein (red), and nuclei with Hoechst (blue). Representative immunofluorescence images show lipid droplet distribution in mock- and HCV-JFH1-infected cells. Lipid droplet fluorescence intensity was quantified on a per-cell basis using ImageJ and compared between HCV core-positive infected cells and matched mock-infected controls. Data are shown as mean ± SEM. n = 600, 720, 745, 725, and 810 cells for 24, 36, 48, 72, and 96 hpi, respectively. Scale bar, 10 µm. Statistical analyses are described in Methods. ns, not significant; ****p < 0.0001.

These findings demonstrate that HCV infection induces dynamic, time-dependent remodeling of LDs. The early increase in LD intensity may reflect enhanced LD biogenesis, expansion, or redistribution during the initial stages of infection. The subsequent decline may indicate that LDs are progressively mobilized or consumed to support ongoing viral replication and particle production. Thus, HCV infection produces a transient increase in LD abundance early after infection, followed by a relative loss of LDs as infection progresses.

### 2. HCV infection induces time-dependent changes in lipid droplet-associated metabolic gene and protein expression

To determine whether the dynamic changes in LDs during HCV infection are associated with remodeling of LD-related metabolic pathways, we performed RNA-seq analysis of mock- and HCV-infected cells at 24 and 48 hpi. HCV infection induced broad transcriptional changes at both time points, with a stronger response at 48 hpi (Figure 2A). Principal component analysis (PCA) showed clear separation between mock and infected samples, while read count distributions were comparable among replicates, supporting the quality and reproducibility of the dataset (Figure 2B–C).

**Figure 2.**
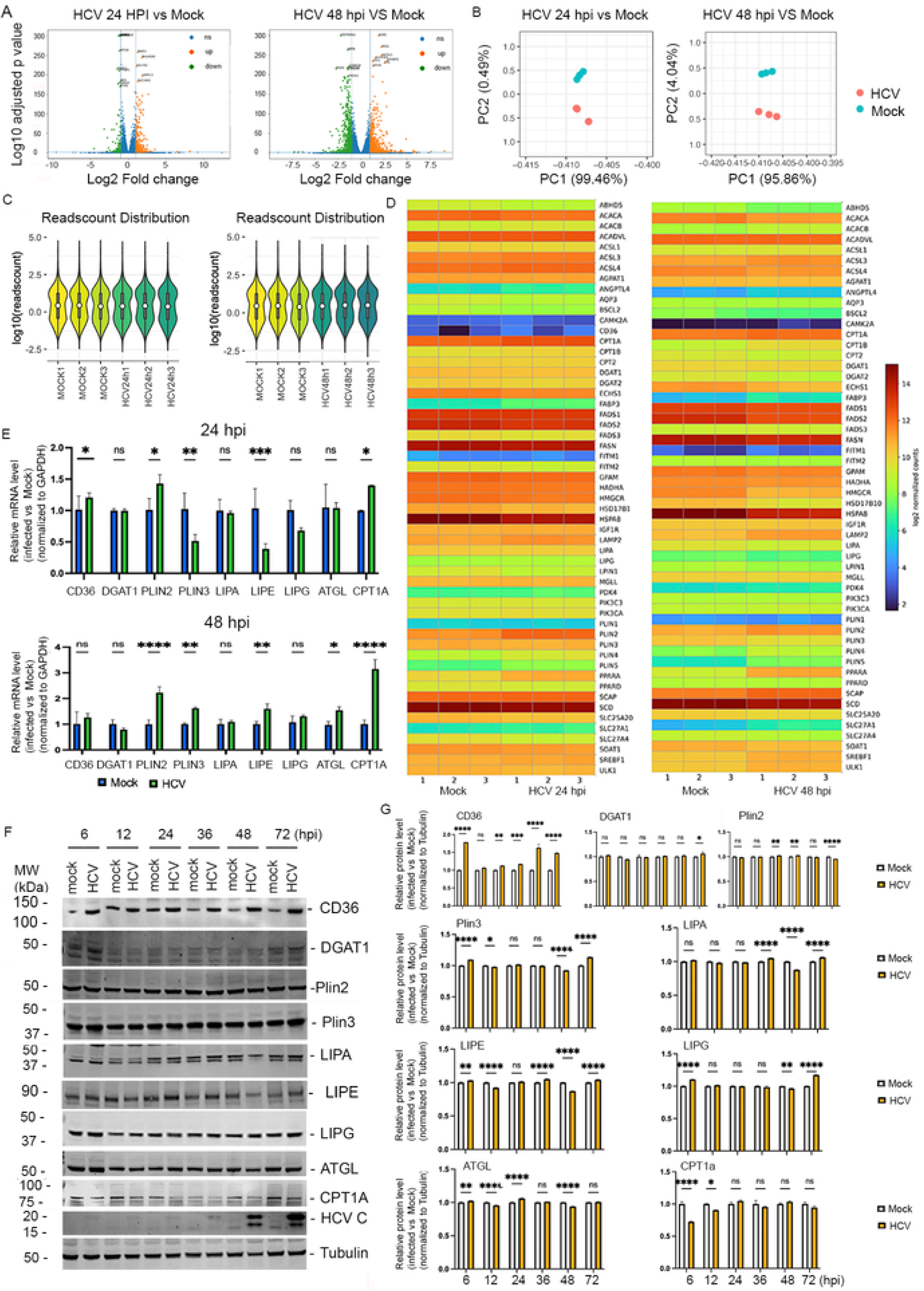
HCV-JFH1 infection induces time-dependent changes in lipid droplet-associated metabolic gene and protein expression. Huh7-derived cells were mock-infected or infected with HCV-JFH1 at an MOI of 1 and harvested at the indicated time points. (A) Volcano plots showing differentially expressed genes in HCV-JFH1-infected cells compared with mock-infected controls at 24 and 48 hpi. Orange indicates upregulated genes, green indicates downregulated genes, and blue indicates genes with no significant change. (B) Principal component analysis of RNA-seq samples. (C) Read count distribution plots showing sequencing read distributions across mock-and HCV-JFH1-infected biological replicates. (D) Heatmaps showing expression patterns of lipid droplet-associated and lipid metabolism-related genes at 24 and 48 hpi. (E) RT-qPCR validation of selected lipid metabolism genes at 24 and 48 hpi. Gene expression was normalized to GAPDH and calculated relative to mock-infected controls using the 2−ΔΔCt method. (F) Western blot analysis of lipid metabolism-associated proteins and HCV core protein across a 6–72 hpi time course. Tubulin was used as a loading control. (G) Quantification of protein expression normalized to tubulin. Data are shown as mean ± SEM. Statistical analyses are described in Methods. ns, not significant; *p < 0.05; **p < 0.01; ***p < 0.001; ****p < 0.0001.

Because HCV replication and assembly are linked to LDs, we next examined genes involved in lipid uptake and transport (CD36), LD structure and stability (Plin2 and Plin3), triglyceride synthesis and LD synthesis (DGAT1), lipolysis and lysosomal lipid mobilization (LIPA, LIPE, LIPG, and ATGL), and mitochondrial fatty acid utilization (CPT1A). Heatmap analysis showed that HCV infection altered this lipid-associated gene network in a time-dependent manner (Figure 2D). RT-qPCR analysis confirmed significant changes in selected genes. At 24 hpi, PLIN2 and CPT1A were increased, while PLIN3 and LIPE were reduced. By 48 hpi, PLIN2, PLIN3, LIPE, ATGL, and CPT1A were significantly increased, suggesting a transition from early LD remodeling toward later activation of lipid mobilization and fatty acid utilization pathways (Figure 2E).

We then examined whether these transcriptional changes were reflected at the protein level. Western blot analysis showed a progressive increase in HCV core protein over time, consistent with ongoing viral replication (Figure 2F). Quantification showed time-dependent regulation of several LD- and lipid metabolism-associated proteins, although the protein profiles did not always match the mRNA changes. CPT1A protein levels remained largely unchanged, whereas PLIN2, DGAT1, and LIPE showed selective increases at later time points. Interestingly, CD36 protein levels were increased across most of the time points, suggesting increased lipid uptake or intracellular lipid transport during infection (Figure 2G). Proteins involved in lipid mobilization, including LIPA, ATGL, LIPG, and LIPE, also displayed distinct early and late expression patterns, consistent with the time-dependent changes in LD abundance shown in Figure 1.

Together, these results show that HCV infection dynamically remodels LD-associated metabolism at both the transcriptional and protein levels. Early infection is characterized predominantly by changes in LD structural and regulatory factors, while later infection is associated with increased expression of lipolytic and fatty acid utilization genes. These results support a model in which HCV progressively redirects host LD metabolism to coordinate LD remodeling, lipid mobilization, and the metabolic demands of viral replication and particle production.

### 3. HCV infection induces time-dependent lipid droplet fusion and enlargement

To determine whether HCV infection alters LD morphology over time, HCV-infected Huh7 cells were stained for LDs, HCV core protein, and nuclei from 24 to 120 hours post-infection (hpi). At 24 hpi, infected cells contained numerous small LDs distributed throughout the cytoplasm, with a subset located near HCV core-positive regions. At 36 and 48 hpi, LDs became more heterogeneous in size and increasingly accumulated in clusters within infected cells. This morphological remodeling became more prominent at later stages of infection. At 72 hpi, enlarged LDs and closely associated LD aggregates were readily detected, whereas by 96 and 120 hpi, infected cells contained markedly enlarged, rounded, or irregularly shaped LD structures (Figure 3A).

**Figure 3.**
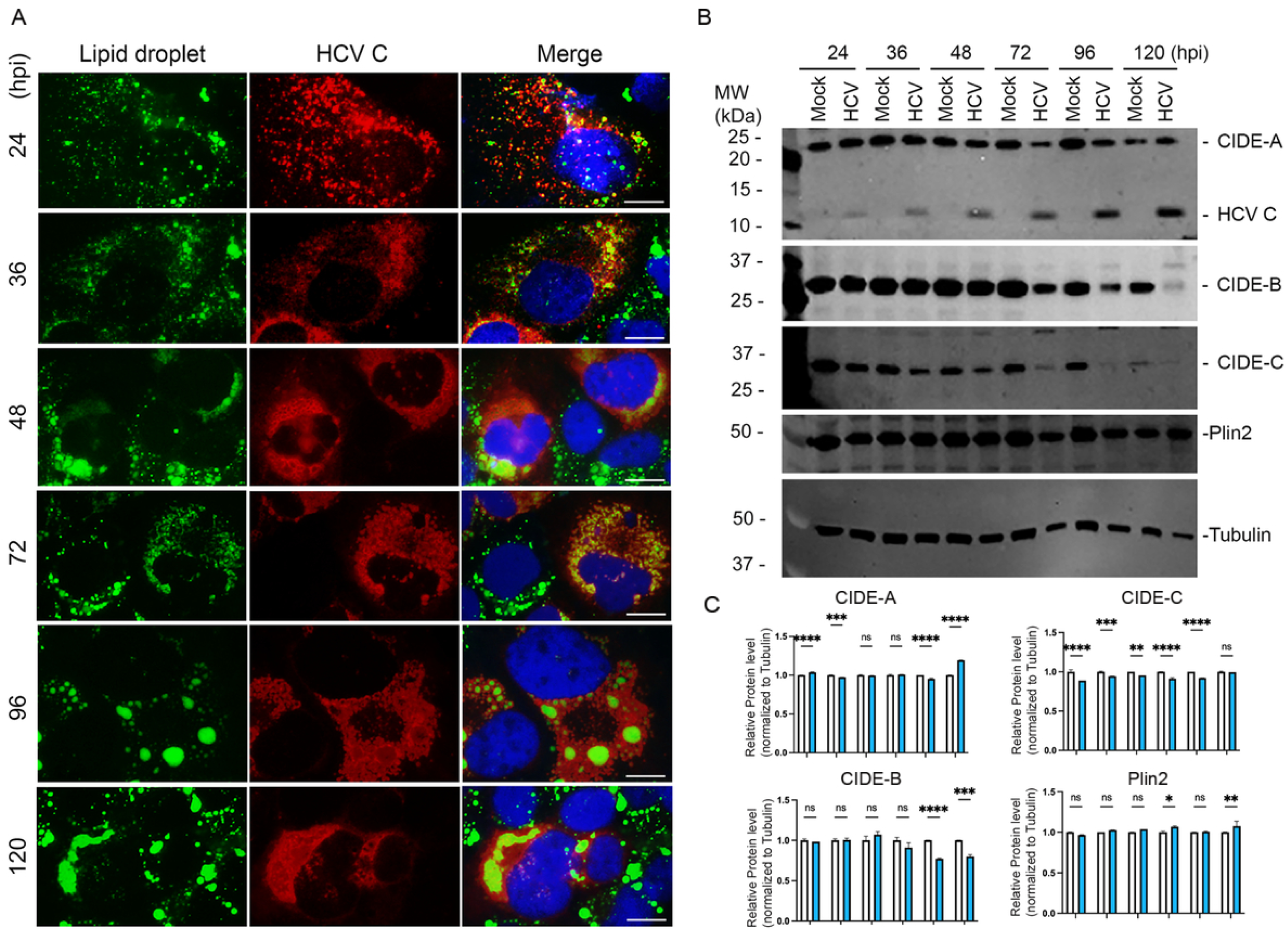
HCV-JFH1 infection is associated with progressive lipid droplet fusion and enlargement despite reduced CIDE protein expression. Huh7 cells were mock-infected or infected with HCV-JFH1 at an MOI of 1 and analyzed at 24, 36, 48, 72, 96, and 120 h post-infection (hpi). **(A)** Representative immunofluorescence images of HCV-JFH1-infected cells stained for lipid droplets using BODIPY 493/503 (green), HCV core protein (red), and nuclei with Hoechst (blue). Merged images show lipid droplet distribution relative to HCV core-positive cells over the course of infection. **(B)** Western blot analysis of CIDEA, CIDEB, CIDEC, PLIN2, and HCV core protein in mock- and HCV-JFH1-infected cells at the indicated time points. Tubulin was used as a loading control. **(C)** Quantification of CIDEA, CIDEB, CIDEC, and PLIN2 protein levels normalized to tubulin. Data are shown as mean ± SEM. Scale bar, 10 µm. Statistical analyses are described in Methods. ns, not significant; *p < 0.05; **p < 0.01; ***p < 0.001; ****p < 0.0001.

The progressive appearance of larger LDs, together with the reduction in the relative abundance of small, dispersed droplets, is consistent with time-dependent LD coalescence or fusion during HCV infection. Enlarged LDs were frequently observed within or adjacent to regions containing abundant HCV core protein, supporting a close spatial relationship between viral infection and LD remodeling. At 120 hpi, some LD structures appeared elongated or interconnected, further suggesting fusion of neighboring droplets or extensive reorganization of LD-associated neutral lipids.

Several LD structural proteins are related to fusion and enlargement of LDs, including CIDE-A, - B, and -C, and Plin2. Therefore, we performed western blot assay (Figure 3B and 3C) to examine the protein levels over the HCV infection through 120 hpi. Unexpectedly, all these LD fusion-related proteins are down-regulated significantly, except Plin2. CIDE-A and -C protein levels are reduced from 24 hpi to 120 hpi, while CIDE-B is down-regulated only during the later stage of infection from 72 hpi. These data suggest that HCV infection-induced LD enlargement may be CIDE-independent.

Together, these observations demonstrate that HCV infection induces progressive changes in LD morphology. Early infection is characterized primarily by numerous small, dispersed LDs, whereas later infection is associated with LD clustering, fusion, and enlargement. These findings suggest that HCV not only alters LD abundance but also reorganizes LD architecture as infection progresses, potentially generating enlarged lipid-storage compartments that contribute to viral replication, assembly, or lipid mobilization.

### 4. HCV infection promotes mitochondrial fatty acid accumulation, increased cellular respiration, and oxidative stress

To determine whether HCV infection alters intracellular fatty acid distribution and mitochondrial metabolism, mock and HCV-infected cells were fractionated into whole-cell lysate, cytosolic, and mitochondrial fractions at 24, 48, and 72 hpi. Western blot analysis confirmed successful subcellular fractionation, with Tom70 enriched in mitochondrial fractions and tubulin detected mainly in whole-cell lysate and cytosolic fractions (Figure 4A). HCV core protein was detected in infected cells and was enriched in mitochondrial fractions, indicating that HCV core associates with mitochondria during infection.

**Figure 4.**
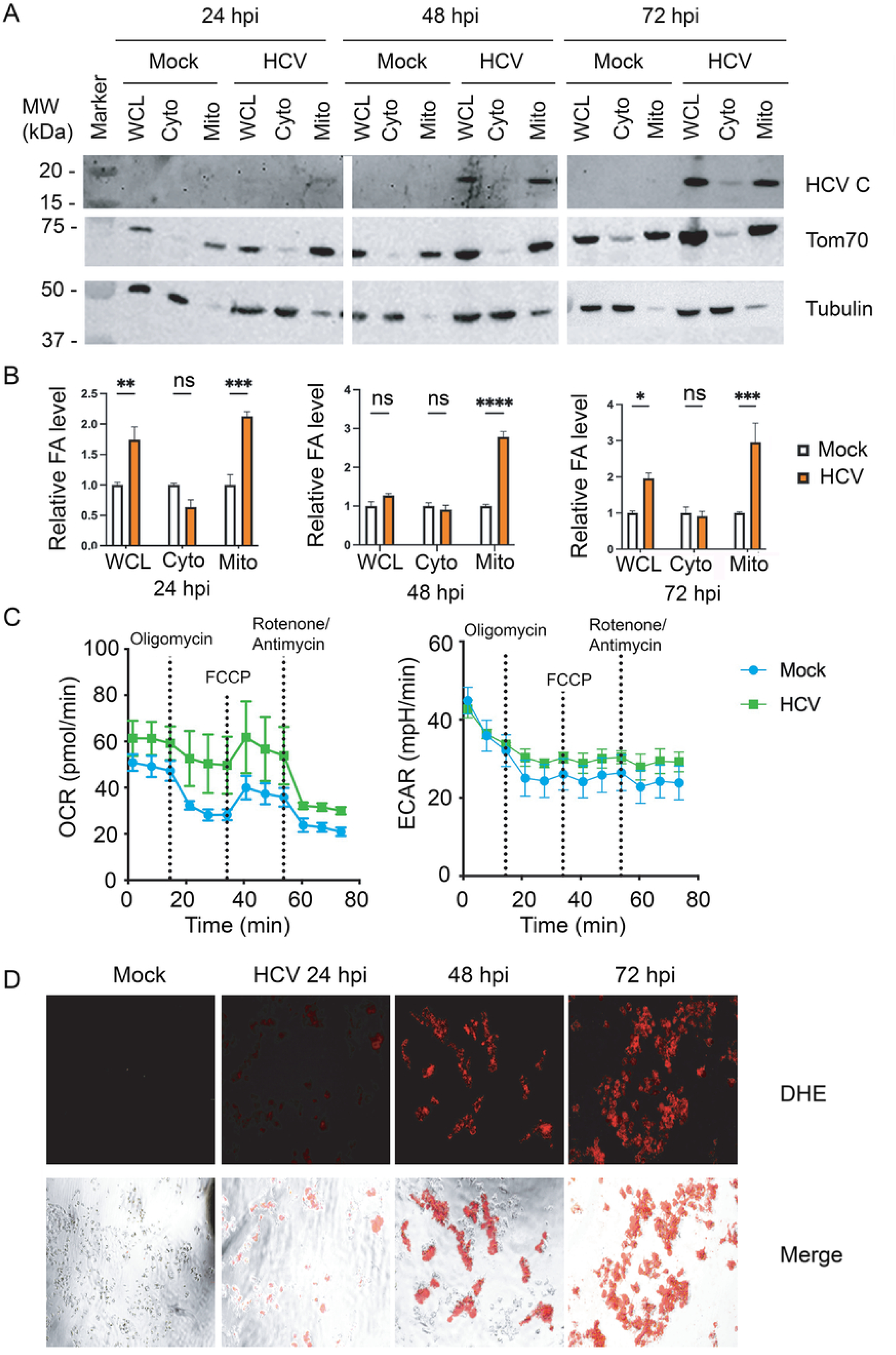
HCV-JFH1 infection promotes mitochondrial fatty acid accumulation, increased respiration, and oxidative stress. Huh7 cells were mock-infected or infected with HCV-JFH1 at an MOI of 1 and harvested at 24, 48, and 72 hpi for subcellular fractionation and fatty acid analysis. (A) Western blot analysis of whole-cell lysate (WCL), cytosolic (Cyto), and mitochondrial (Mito) fractions from mock- and HCV-JFH1-infected cells. HCV core protein was used to confirm infection, Tom70 was used as a mitochondrial marker, and tubulin was used as a cytosolic and loading control. (B) Free fatty acids were quantified using a colorimetric free fatty acid assay in WCL, cytosolic, and mitochondrial fractions and normalized to matched mock-infected controls. (C) Seahorse XF Cell Mito Stress Test analysis of oxygen consumption rate (OCR) and extracellular acidification rate (ECAR) in mock- and HCV-JFH1-infected Huh7.0 cells at 48 hpi. For Seahorse analysis, cells were infected with HCV-JFH1 at an MOI of 3 and analyzed following sequential injection of oligomycin, FCCP, and rotenone/antimycin A. (D) Reactive oxygen species levels were measured in mock- and HCV-JFH1-infected Huh7.0 cells at 24, 48, and 72 hpi. ROS was quantified by fluorescence intensity relative to mock controls. Data are shown as mean ± SEM. Statistical analyses are described in Methods. ns, not significant; *p < 0.05; **p < 0.01; ***p < 0.001; ****p < 0.0001.

We next measured fatty acid levels in each subcellular fraction. As shown in Figure 4B, HCV infection significantly increased total cellular fatty acid levels (whole-cell lysate subfraction) at 24 and 72 hpi, whereas cytosolic fatty acid levels were not significantly changed. In contrast, mitochondrial fatty acid levels were significantly increased at 24, 48, and 72 hpi. These findings indicate that HCV infection promotes preferential accumulation or trafficking of fatty acids to mitochondria rather than their general accumulation in the cytosol.

To assess whether mitochondrial fatty acid accumulation was associated with altered mitochondrial metabolism, we measured oxygen consumption rate (OCR) and extracellular acidification rate (ECAR) in mock and HCV-infected (48 hpi) Huh7 cells. HCV-infected cells showed an increased OCR compared with mock-infected controls, indicating enhanced cellular respiration. ECAR was also modestly elevated, suggesting that HCV infection concurrently increases extracellular acidification and may enhance glycolytic activity (Figure 4C).

Finally, intracellular reactive oxygen species were assessed by 2′,7′-dichlorofluorescin diacetate (DCFDA)-based ROS detection Assay staining. exhibited increased red fluorescence compared with mock-infected controls, and the signal progressively increased from 24 to 72 hpi. These results demonstrate that HCV infection induces time-dependent accumulation of reactive oxygen species and oxidative stress (Figure 4D).

Together, these findings show that HCV infection promotes the redistribution of fatty acids to mitochondria, increases cellular respiration, and induces progressive oxidative stress. These metabolic changes support a model in which HCV mobilizes host fatty acids for mitochondrial utilization while simultaneously increasing the oxidative burden of infected cells.

### 5. Inhibition of LD biogenesis, lipolysis, and mitochondrial fatty acid transport reduces HCV-induced mitochondrial fatty acid accumulation and metabolic activation

To determine whether HCV-induced mitochondrial fatty acid accumulation depends on LD biogenesis, lipid hydrolysis, or mitochondrial fatty acid transport, HCV-infected cells were treated with Etomoxir, an inhibitor of CPT1; A922500 (DGAT1 inhibitor), Lalistat 2 (lysosomal acid lipase/LIPA inhibitor), or HSL-In-1 (hormone-sensitive lipase/LIPE inhibitor). Cells were analyzed at 48 hpi. Cell viability assays were performed using MTT (3-[4,5-dimethylthiazol-2-yl]-2,5 diphenyl tetrazolium bromide) assay to identify concentrations below the approximate CC50 values, which were 1.0 µM for Etomoxir, 8.0 µM for A922500, 25 µM for Lalistat 2, and 10 µM for HSL-In-1. Based on these results, cells were treated with 0.5 µM Etomoxir, 2.5 µM A922500, 2.5 µM Lalistat 2, or 2.5 µM HSL-In-1 (Figure 5A).

**Figure 5.**
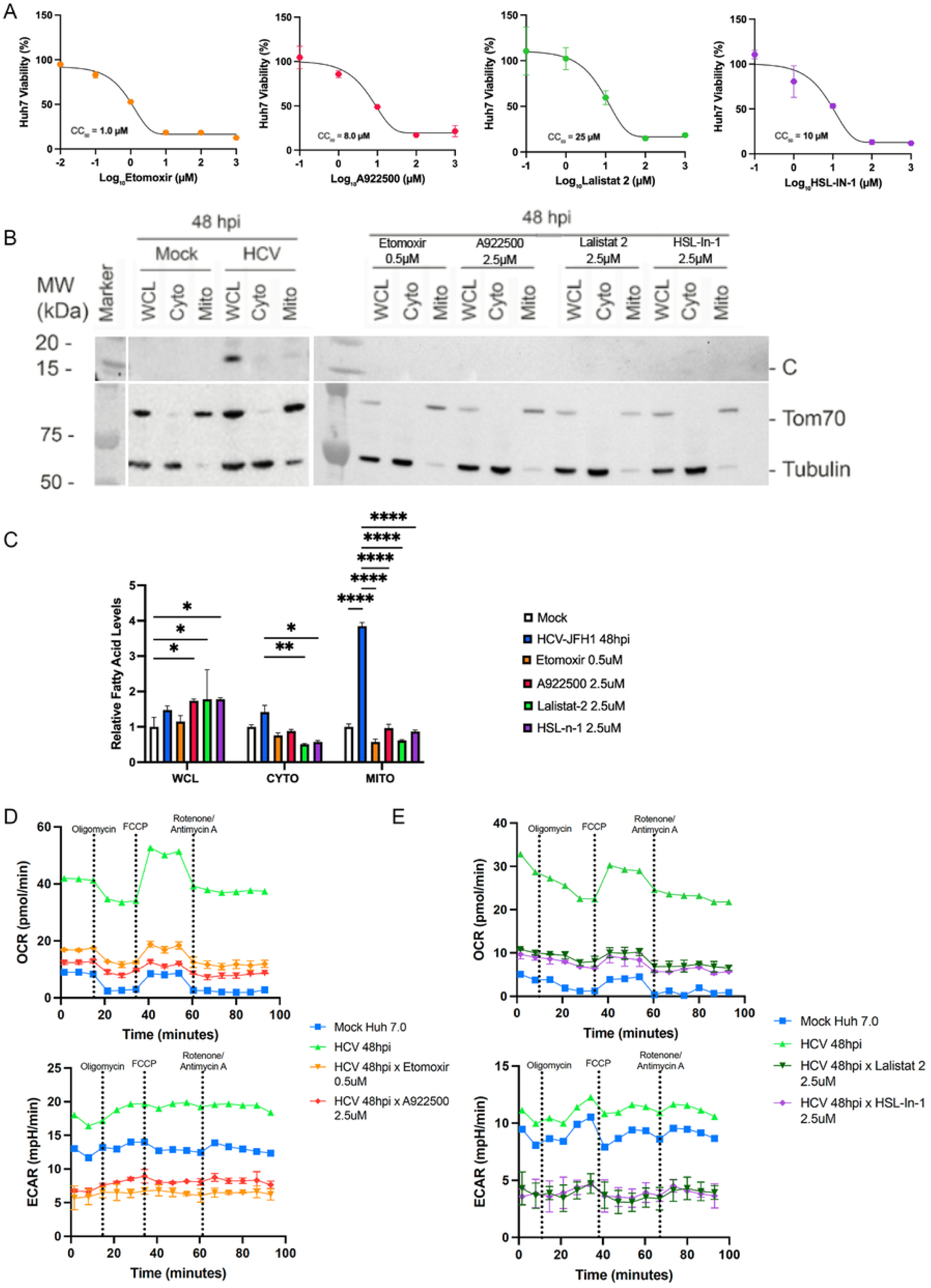
**Inhibition of fatty acid transport and lipolysis reduces HCV-JFH1-inducedmitochondrial fatty acid accumulation and metabolic activation.**Huh7 cells were mock-infected or infected with HCV-JFH1 at an MOI of 1 and treated with Etomoxir, A922500, Lalistat 2, or HSL-In-1 as indicated. **(A)** Cell viability was measured by MTT assay following inhibitor treatment. CC₅₀ values were approximately 1.0 µM for Etomoxir, 8.0 µM for A922500, 25 µM for Lalistat 2, and 10 µM for HSL-In-1. **(B)** Western blot analysis of mock- and HCV-JFH1-infected cells treated with Etomoxir, A922500, Lalistat 2, or HSL-In-1 at 48 hpi. Cells were fractionated into whole-cell lysate (WCL), cytosolic (Cyto), and mitochondrial (Mito) fractions. HCV core protein was used to confirm infection, Tom70 was used as a mitochondrial marker, and tubulin was used as a cytosolic and loading control. **(C)** Free fatty acids were quantified using a colorimetric free fatty acid assay in WCL, cytosolic, and mitochondrial fractions and normalized to matched controls. **(D)** Seahorse XF Cell Mito Stress Test analysis of oxygen consumption rate (OCR) and extracellular acidification rate (ECAR) in mock- and HCV-JFH1-infected Huh7.0 cells treated with Etomoxir or A922500. **(E)** Seahorse XF Cell Mito Stress Test analysis of OCR and ECAR in mock- and HCV-JFH1-infected Huh7.0 cells treated with Lalistat 2 or HSL-In-1. For Seahorse analysis, cells were infected with HCV-JFH1 at an MOI of 3 and analyzed at 48 hpi following sequential injection of oligomycin, FCCP, and rotenone/antimycin A. Data are shown as mean ± SEM. Statistical analyses are described in Methods. ns, not significant; *p < 0.05; **p < 0.01; ****p < 0.0001.

Whole-cell lysate, cytosolic, and mitochondrial fractions were isolated. Western blot analysis confirmed the successful subcellular fractionation, with Tom70 enriched in mitochondrial fractions and tubulin detected mainly in whole-cell lysate and cytosolic fractions. HCV core protein was detected in infected cells, confirming infection (Figure 5B).

HCV infection increased fatty acid abundance in all three fractions at 48 hpi, with the strongest increase observed in the mitochondrial fraction. Relative to mock-infected cells, HCV infection increased fatty acid levels by approximately 1.5-fold in whole-cell lysate, 1.4-fold in the cytosolic fraction, and 3.5- to 4-fold in the mitochondrial fraction. Inhibitor treatment strongly reduced the mitochondrial fatty acid pool. Etomoxir, A922500, Lalistat 2, and HSL-In-1 decreased mitochondrial fatty acid levels from approximately 3.8-fold in untreated HCV-infected cells to near baseline levels, representing an approximately 70–85% reduction (Figure 5C).

In contrast, the effects of these inhibitors on whole-cell lysate and cytosolic fractions were less uniform. Whole-cell fatty acid levels remained similar to or slightly higher than those in untreated HCV-infected cells, while cytosolic fatty acid levels were reduced modestly, particularly after Lalistat 2 and HSL-In-1 treatment. These findings suggest that inhibition of these pathways primarily disrupts fatty acid mobilization and delivery to mitochondria rather than uniformly reducing the total cellular fatty acid pool. The results further indicate that DGAT1-dependent LD biogenesis, LIPA-dependent lysosomal lipid hydrolysis, LIPE/HSL-dependent lipolysis, and CPT1-dependent mitochondrial fatty acid transport all contribute to the mitochondrial fatty acid accumulation induced by HCV infection.

We next measured OCR and ECAR to determine whether the reduction in mitochondrial fatty acid accumulation was associated with altered metabolic activity. HCV infection increased OCR approximately 4 to 5-fold compared with mock-infected cells, indicating marked activation of cellular respiration. Etomoxir or A922500 reduced OCR to approximately 25–40% of the level observed in untreated HCV-infected cells. Similarly, Lalistat 2 and HSL-In-1 reduced OCR close to levels approaching those of mock-infected controls, indicating that both lysosomal lipid hydrolysis and HSL-dependent lipolysis contribute to HCV-induced respiratory activation (Figure 5D).

HCV infection also increased ECAR compared with mock-infected cells, suggesting increased extracellular acidification. Inhibitor treatment reduced ECAR, with Lalistat 2 and HSL-In-1 producing the strongest suppression. Thus, disruption of fatty acid mobilization and mitochondrial import attenuated both respiratory and extracellular acidification responses during HCV infection (Figure 5E).

Together, these results demonstrate that HCV-induced metabolic activation depends on coordinated LD biogenesis, lipolytic fatty acid mobilization, and mitochondrial fatty acid transport. Inhibition of DGAT1, LIPA, LIPE/HSL, or CPT1 markedly reduced mitochondrial fatty acid accumulation and suppressed the increases in OCR and ECAR associated with HCV infection. These findings support a model in which HCV mobilizes fatty acids from LD-associated lipid pools and directs them to mitochondria to sustain infection-associated metabolic activity.

### 6. Inhibition of lipid metabolic pathways differentially reduces HCV RNA and core protein levels

To determine whether LD-associated metabolic pathways contribute to HCV replication and viral protein accumulation, HCV-infected cells were treated with Etomoxir (CPT1 inhibitor), A922500 (DGAT1 inhibitor), Lalistat 2 (lysosomal acid lipase/LIPA inhibitor), or HSL-In-1 (hormone-sensitive lipase/LIPE inhibitor), HCV RNA and core protein levels were analyzed at 48 hpi. RT-qPCR analysis showed that inhibitors had distinct effects on HCV RNA abundance (Figure 6A). Etomoxir did not significantly alter HCV RNA levels at any of the concentrations tested, suggesting that inhibition of CPT1-dependent mitochondrial fatty acid transport had little effect on viral RNA abundance under these conditions. In contrast, A922500 reduced HCV RNA levels by approximately 20–40%, with the strongest reduction observed at the highest dose. Lalistat 2 also reduced HCV RNA in a dose-dependent manner, with viral RNA decreasing by approximately 40–50% at higher concentrations. HSL-In-1 produced a similar reduction, lowering HCV RNA by approximately 30–50% compared with untreated HCV-infected cells. These findings suggest that DGAT1-dependent lipid metabolism, LIPA-mediated lysosomal lipid hydrolysis, and LIPE/HSL-dependent lipolysis contribute to efficient HCV RNA accumulation, whereas CPT1 inhibition has a comparatively limited effect on viral RNA.

**Figure 6.**
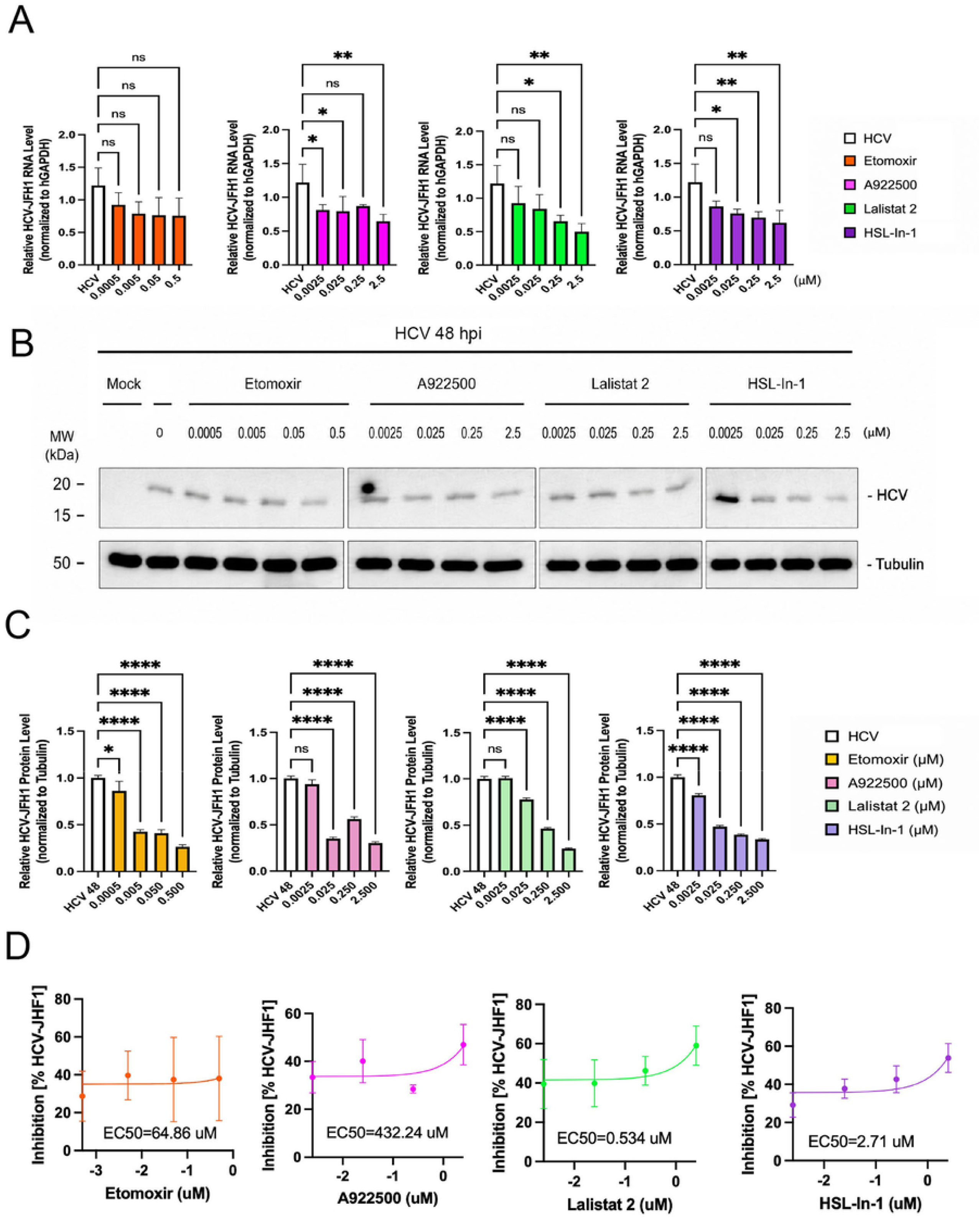
Inhibition of lipid metabolic pathways differentially reduces HCV-JFH1 RNA and core protein levels. Huh7 cells were infected with HCV-JFH1 at an MOI of 1 and treated with Etomoxir, A922500, Lalistat 2, or HSL-In-1 as indicated. RNA and protein were harvested at 48 hpi following inhibitor treatment. (A) RT-qPCR analysis of HCV-JFH1 RNA levels in inhibitor-treated infected cells. Viral RNA levels were normalized to GAPDH and calculated relative to untreated HCV-JFH1-infected controls using the 2−ΔΔCt method. (B) Western blot analysis of HCV core protein in inhibitor-treated cells at 48 hpi. Tubulin was used as a loading control. Quantification of HCV core protein levels normalized to tubulin. (D) Dose-response curves showing percent inhibition of HCV-JFH1 following inhibitor treatment. Estimated EC50 values were 64.86 µM for Etomoxir, 432.24 µM for A922500, 0.534 µM for Lalistat 2, and 2.71 µM for HSL-In-1. Data are shown as mean ± SEM. Statistical analyses are described in Methods. ns, not significant; *p < 0.05; **p < 0.01; ****p < 0.0001.

We next examined HCV core protein levels by western blot (Figure 6B). All four inhibitors reduced HCV core protein accumulation, although their concentration-response patterns differed. Etomoxir reduced HCV core protein levels by approximately 60–75%, despite having minimal effect on viral RNA. A922500 also decreased HCV core protein, with the strongest reduction observed at the highest concentration. Lalistat 2 reduced HCV core protein in a dose-dependent manner, with protein levels reduced by approximately 70–80% at the highest dose. HSL-In-1 similarly decreased HCV core protein levels by approximately 60–70% (Figure 6C). These results suggest that inhibition of DGAT1-dependent lipid droplet biogenesis, LIPA-dependent lysosomal lipid hydrolysis, LIPE/HSL-dependent lipolysis, and CPT1-dependent fatty acid transport can impair viral protein accumulation, although their effects on viral RNA are not equivalent.

Notably, etomoxir markedly reduced HCV core protein despite having no significant effect on viral RNA. A922500 also produced a greater reduction in core protein than in viral RNA. These differences indicate that CPT1- and DGAT1-dependent pathways may have stronger effects on post-RNA-replication events, such as viral protein synthesis or stability, LD-associated assembly, or virion production, than on viral RNA accumulation itself. In contrast, inhibition of LIPA or LIPE/HSL reduced both viral RNA and core protein, suggesting broader roles for lipid hydrolysis and fatty acid mobilization in the HCV life cycle.

Dose-response analysis further showed that Lalistat 2 and HSL-In-1 had stronger inhibitory activity against HCV than Etomoxir or A922500 (Figure 6D). The estimated EC50 values were approximately 0.534 µM for Lalistat 2 and 2.71 µM for HSL-In-1, compared with 64.86 µM for Etomoxir and 432.24 µM for A922500. Together, these data suggest that lipid hydrolysis pathways, particularly LIPA-dependent lysosomal lipid hydrolysis and LIPE/HSL-dependent lipolysis, contribute to HCV replication and viral protein production. The stronger reduction in HCV core protein compared with viral RNA suggests that these pathways may affect post-replication steps such as viral protein accumulation, lipid droplet-associated assembly, or particle production.

Together, these results demonstrate that disruption of LD-associated lipid metabolism impairs HCV infection at multiple stages. LIPA-dependent lysosomal lipid hydrolysis and LIPE/HSL-dependent lipolysis support both viral RNA accumulation and core protein production, whereas CPT1-dependent mitochondrial fatty acid transport and DGAT1-dependent lipid metabolism exert more pronounced effects on viral protein accumulation than on viral RNA. These findings support a model in which lipid mobilization and trafficking contribute not only to HCV RNA replication but also to downstream processes required for viral protein accumulation, assembly, and particle production.

## DISCUSSION

HCV has a well-established dependence on host LDs for viral assembly and particle production^32, 41, 42^, but the temporal relationship among LD remodeling, lipid mobilization, mitochondrial metabolism, and viral replication has remained incompletely defined. In this study, we show that HCV infection induces a coordinated and time-dependent reorganization of host lipid metabolism. Early infection was characterized by a transient increase in LD intensity, whereas later infection was associated with progressive LD clustering and enlargement, increased expression of selected lipolytic and fatty acid utilization pathways, preferential accumulation of fatty acids in mitochondria, enhanced cellular respiration, and oxidative stress. Pharmacological disruption of LD biogenesis, lipid hydrolysis, or mitochondrial fatty acid transport reduced mitochondrial fatty acid accumulation and metabolic activation and impaired HCV RNA or core protein accumulation. Together, these findings support a model in which HCV infection first expands or reorganizes LD-associated lipid stores and later promotes the redistribution of fatty acids toward mitochondria, where they may contribute to infection-associated metabolic activation and biosynthetic demands.

The transient increase in LD intensity at 24 hpi is consistent with the established role of LDs during the early stages of the HCV life cycle. HCV core protein localizes to LD surfaces, while NS5A and viral replication complexes are recruited to adjacent ER membranes, thereby bringing newly synthesized viral RNA into proximity with capsid assembly sites^41, 43^. Increased LD abundance early after infection could therefore increase the availability of membrane-associated platforms needed to coordinate replication and assembly. This early increase may result from enhanced triglyceride synthesis, LD biogenesis, stabilization of LDs by perilipins, or redistribution of pre-existing neutral lipids. The accompanying increase in PLIN2 and CPT1A transcripts at 24 hpi indicates that LD structural remodeling and fatty acid utilization pathways are activated early during infection.

At later time points, LDs became more heterogeneous, clustered, and enlarged despite unchanged total LD signal, a pattern that could reflect neutral-lipid redistribution, selective LD growth, preferential utilization of smaller droplets, or coalescence of closely apposed LDs. Importantly, reduced total LD signal and increased apparent LD size are not mutually exclusive, because selective loss of small droplets, clustering of adjacent droplets, or redistribution of neutral lipids into fewer larger structures could reduce overall LD abundance while increasing the size of remaining LD-positive structures. A notable finding was that CIDEA, CIDEB, and CIDEC protein levels declined during the period in which LD enlargement became most evident. Because CIDE proteins are commonly associated with LD growth and fusion, their reduction argues against a simple CIDE-driven mechanism. Instead, HCV-induced LD enlargement may occur through CIDE-independent processes, including DGAT1-dependent triglyceride synthesis, ER-to-LD lipid transfer, HCV core-induced clustering, or PLIN2-mediated stabilization.

Together, these findings suggest that HCV may uncouple LD enlargement from lipid storage. Reduced CIDE protein levels could increase triglyceride accessibility to lipases, allowing enlarged LDs to function as dynamically remodeled lipid reservoirs that support viral assembly and fatty acid mobilization. Future live-cell imaging or lipid-transfer assays will be needed to distinguish true fusion from growth, clustering, or selective loss of smaller droplets.

The transcriptional and protein analyses further support a temporal shift from LD remodeling toward lipid mobilization. HCV altered genes involved in LD organization, triglyceride synthesis, lipid uptake, lipolysis, and mitochondrial fatty acid utilization. At 24 hpi, increased PLIN2 and CPT1A and reduced PLIN3 and LIPE suggested an early phase dominated by LD reorganization and selective regulation of fatty acid handling. By 48 hpi, PLIN2, PLIN3, LIPE, ATGL, and CPT1A transcripts were increased, indicating broader activation of pathways involved in LD stabilization, lipolysis, and mitochondrial fatty acid utilization. The incomplete correspondence between mRNA and protein levels is not unexpected, because lipid metabolic enzymes are also controlled by translation, protein stability, phosphorylation, intracellular localization, and interactions with LD-associated cofactors. These differences emphasize the importance of integrating transcriptional measurements with functional metabolic assays.

Subcellular fractionation revealed that mitochondrial fatty acid levels increased at all examined time points, whereas cytosolic fatty acid abundance changed relatively little. This preferential enrichment suggests that HCV does not merely increase the total cellular fatty acid pool but actively promotes fatty acid delivery to, retention within, or association with mitochondria. The detection of HCV core protein in mitochondrial fractions is also consistent with direct or indirect interactions between viral proteins and mitochondrial membranes. However, biochemical fractionation alone cannot distinguish whether core protein is imported into mitochondria, associates with the outer mitochondrial membrane, or remains bound to mitochondria-associated ER or LD membranes. Definitive localization will require higher-resolution imaging and purification of mitochondria-associated membrane fractions.

The increase in mitochondrial fatty acids was accompanied by elevated oxygen consumption, suggesting that HCV infection enhances respiratory activity under the conditions used in this study. The concurrent increase in extracellular acidification indicates that HCV may stimulate both oxidative and glycolytic metabolism rather than producing a simple switch between these pathways. Such metabolic flexibility may help infected cells meet the increased demands for ATP, membrane synthesis, protein production, and virion assembly. At the same time, enhanced respiration was associated with progressive ROS accumulation, indicating that metabolic activation imposes an oxidative burden. This finding may be relevant to HCV-associated liver injury, because persistent mitochondrial dysfunction and oxidative stress can contribute to lipid peroxidation, inflammatory signaling, hepatocyte damage, fibrosis, and carcinogenesis.

Previous studies have reported that HCV can impair mitochondrial β-oxidation or suppress components of fatty acid oxidation machinery. Our findings do not necessarily conflict with those observations. HCV-induced metabolic effects may vary according to viral genotype, cell type, infection stage, nutritional conditions, and the specific mitochondrial parameters measured. Increased mitochondrial fatty acid accumulation and OCR do not prove that all incoming fatty acids undergo complete β-oxidation. Fatty acid accumulation may reflect delivery to mitochondria that exceeds oxidative capacity, incorporation into mitochondrial membrane lipids, or partial impairment of β-oxidation. The increase in ROS is compatible with a state in which mitochondrial substrate delivery and electron transport are enhanced but incompletely coupled. Direct measurements of fatty acid oxidation, respiratory-chain function, and isotope-labeled lipid flux will be needed to define the fate of this mitochondrial fatty acid pool.

The inhibitor studies provide functional evidence that LD biogenesis, lipid hydrolysis, and mitochondrial fatty acid transport cooperate to generate the HCV-induced mitochondrial fatty acid phenotype. Inhibition of DGAT1, LIPA, LIPE/HSL, or CPT1 markedly reduced mitochondrial fatty acid accumulation while producing less uniform effects on whole-cell and cytosolic fatty acid levels. These findings suggest that the inhibitors primarily disrupt fatty acid trafficking and compartmentalization rather than simply depleting total cellular fatty acids. The reduction in OCR and ECAR further indicates that LD-derived lipid mobilization is linked to infection-associated metabolic activation.

Together, these inhibitor studies suggest that HCV-induced lipid remodeling requires coordinated triglyceride synthesis, LD turnover, and mitochondrial fatty acid utilization. The effect of A922500 indicates that DGAT1-dependent triglyceride synthesis and LD biogenesis contribute to the formation of a lipid pool that can subsequently be mobilized during infection. Although DGAT1 promotes neutral-lipid storage, newly formed LDs may function as transient metabolic intermediates that collect and organize fatty acids before delivering them to viral assembly sites or mitochondria. Thus, LD biogenesis and LD breakdown should not be viewed as opposing processes, but rather as linked steps in an HCV-induced lipid cycle in which fatty acids are esterified into triglycerides, packaged into LDs, and later released through lysosomal hydrolysis or cytosolic lipolysis. Consistent with this model, Lalistat 2 and HSL-IN-1 identify LIPA-dependent lysosomal hydrolysis of cholesteryl esters and triglycerides and LIPE/HSL-dependent mobilization of cytosolic neutral-lipid stores as distinct but complementary pathways that contribute to mitochondrial fatty acid accumulation, respiratory activation, HCV RNA accumulation, and core protein expression. In contrast, etomoxir produced a distinct phenotype, with minimal effect on HCV RNA but marked reduction of core protein accumulation and HCV-induced respiration, suggesting that CPT1-dependent mitochondrial fatty acid transport is more important for mitochondrial metabolic activation and post-replicative stages of infection than for maintenance of viral RNA abundance. A similar, although less pronounced, divergence after DGAT1 inhibition further suggests that viral RNA and protein measurements reflect different stages of the viral life cycle and that host lipid pathways may support HCV infection at multiple mechanistic points.

The apparent antiviral potencies of the inhibitors should be interpreted with appropriate caution. Lalistat 2 and HSL-IN-1 produced measurable concentration-dependent inhibition within the tested ranges, whereas the calculated EC50 values for etomoxir and A922500 exceeded the concentrations evaluated. Those extrapolated values therefore do not provide precise estimates of antiviral potency. In addition, pharmacological inhibitors can have off-target or pathway-independent effects, particularly at higher concentrations. Future studies should combine inhibitor experiments with siRNA, CRISPR-mediated depletion, rescue with inhibitor-resistant proteins, and direct measurements of viral infectivity to validate the specific contributions of CPT1A, DGAT1, LIPA, and LIPE.

Several limitations should be considered. This study was performed primarily in Huh7-derived cells infected with genotype 2a JFH-1, and responses may differ in primary hepatocytes, liver organoids, or other HCV genotypes. Because extended HCV infection may be associated with cytopathic stress or reduced cell viability at later time points, future studies should pair LD and mitochondrial measurements with direct viability, apoptosis, or cell death assays across the infection time course. Fatty acid measurements in biochemical fractions did not identify specific lipid species or distinguish free from lipid-associated fatty acids. Increased OCR suggests enhanced respiration, but detailed respiratory profiling is needed to determine whether mitochondrial activity is productive or dysfunctional. Finally, infectious virion production was not directly measured, and inhibitor effects on core protein should be tested using infectivity assays.

Despite these limitations, the study provides an integrated model for temporal LD remodeling during HCV infection. In the proposed model, early infection promotes LD biogenesis, stabilization, or redistribution to establish replication- and assembly-competent membrane domains. As infection progresses, LDs become clustered and enlarged through CIDE-independent mechanisms, while lipolytic and lysosomal pathways mobilize stored fatty acids. These fatty acids are preferentially redirected toward mitochondria, where they support infection-associated respiratory activity but also promote ROS accumulation and oxidative stress. DGAT1-dependent LD biogenesis, LIPA- and LIPE-dependent lipid hydrolysis, and CPT1-dependent mitochondrial fatty acid transport act as interconnected components of this metabolic program.

In conclusion, our findings expand the role of LDs in HCV infection beyond their established function as viral assembly platforms. LDs also appear to serve as dynamic metabolic intermediates that coordinate lipid storage, remodeling, hydrolysis, and mitochondrial substrate delivery. The identification of LIPA- and LIPE/HSL-dependent lipid hydrolysis as important determinants of viral RNA and core protein accumulation highlights these pathways as potential host-directed antiviral targets. Targeting the metabolic interface among LDs, lipolysis, and mitochondria may provide a strategy to inhibit HCV infection while also limiting the oxidative and metabolic disturbances that contribute to HCV-associated liver disease.

## METHODS and MATERIALS

### Cell culture and viral infection

Huh7 cells (a gift from Dr. Yanjin Zhang at University of Maryland) were cultured and maintained in Dulbecco’s Modified Eagle’s Medium (DMEM) supplemented with 10% fetal calf serum (FCS), penicillin–streptomycin (100 IU/mL penicillin and 100 µg/mL streptomycin), and amphotericin B (2.5 µg/mL). HCV was generated from the JFH-1 clone, which was a kind gift from Prof. Takaji Wakita (National Institute of Infectious Diseases, Tokyo, Japan). To generate genomic HCV RNA, plasmid pJFH1 was linearized with XbaI, treated with mung bean nuclease, and used as a template for in vitro transcription with the MEGAscript kit from Ambion. In vitro-transcribed RNA was transfected into Huh7 cells by electroporation as previously described. Cell culture supernatant was collected, and viral titer was determined by immunofluorescence assay using an anti-HCV capsid/core antibody.

### Immunofluorescence assay and lipid droplet staining

Human hepatoma-derived Huh7 cells were seeded on coverslips in 24-well plates at 0.2 × 10⁶ cells per well one day before infection. Cells were infected with HCV JFH-1 and incubated for 12, 24, 48, 72, 96, and 120 h with mock-infected cells included as matched uninfected controls. After the designated infection period, coverslips were fixed with 2% formaldehyde and rinsed three times with PBS. Cells were then permeabilized with 0.2% Triton X-100. Lipid droplets were stained with BODIPY 493/503 dye at a dilution of 1:1000 in MEM for 30 min. To identify infected cells, an anti-HCV core protein antibody was used as a viral marker. Cells were visualized under a fluorescence microscope using matched exposure and magnification across all conditions. ImageJ was used to segment cells and quantify lipid droplet size, number, and fluorescence intensity on a per-cell basis. Analysis was restricted to HCV-positive cells in comparison with matched mock-infected cells. More than three biological replicates were included, with a target of 500–1000 cells analyzed per condition.

### RNA-sequencing data analysis

RNA-sequencing data were analyzed in Python using raw, unnormalized gene-level count matrices. Count data were imported with pandas, cleaned to remove missing or duplicated gene identifiers, converted to numeric format, and filtered to exclude genes with fewer than 10 total counts across the samples in each comparison. HCV-JFH1 infected Huh7 cells at 24 and 48 hours were analyzed separately against their corresponding mock-treated controls, resulting in two independent comparisons: HCV 24 h versus mock and HCV 48 h versus mock. Differential expression analysis was performed with PyDESeq2 using a negative-binomial generalized linear model with treatment condition as the only factor and mock-treated samples as the reference group. Genes were considered differentially expressed when the adjusted P-value was < 0.05 and the absolute log2 fold change was ≥ 1.

For visualization and quality assessment, raw counts were normalized using a DESeq-style median-ratio method and transformed as log2(normalized count + 1). These transformed values were used for principal component analysis, sample-correlation analysis, volcano plots, and heatmaps, while differential expression testing was performed using untransformed integer counts. Functional enrichment analysis was performed with GSEApy and the Enrichr API using upregulated genes with adjusted P-value < 0.10 and log2 fold change ≥ 0.5. Enrichment was assessed against Gene Ontology Biological Process 2023, KEGG Human 2021, and Reactome 2022 gene sets, with targeted interpretation of pathways related to endoplasmic-reticulum stress, unfolded protein response, protein processing in the endoplasmic reticulum, lipid droplets, lipid metabolism, and fatty-acid β-oxidation. All graphical outputs were generated with Matplotlib and exported at 300 dpi.

### MTT (3-[4,5-dimethylthiazol-2-yl]-2,5 diphenyl tetrazolium bromide) assay for cell viability

Cell viability was assessed using the MTT assay. Cells were seeded into 96-well plates in Minimal Essential Medium (MEM) supplemented with 10% fetal bovine serum (FBS) and allowed to attach overnight under standard culture conditions at 37 °C with 5% CO₂. Following treatment, the culture medium was aspirated and replaced with fresh medium containing the appropriate experimental conditions. Cells were incubated for 24, 36, 48, 72, 96, or 120 h at 37 °C. After incubation, cells were gently washed with PBS. MTT solution (0.5 mg/mL; Sigma, lot #MKCD8033) prepared in MEM containing 10% FBS was added to each well, and plates were incubated for 2 h at 37 °C. During this period, metabolically active cells reduced the tetrazolium salt to insoluble purple formazan crystals. Following incubation, the MTT-containing medium was carefully removed, and 100 µL of isopropanol was added to each well to dissolve the formazan crystals. Plates were incubated for 15 min at room temperature with gentle shaking to ensure complete solubilization. Absorbance was measured using a microplate reader spectrophotometer at 570 nm, with 690 nm used as a reference wavelength. Cell viability was calculated based on absorbance values and expressed as a percentage relative to untreated control cells, which were defined as 100% viable. All experiments were performed in at least three independent biological replicates.

### Western blot analysis

Western blot analysis was performed to examine viral and host protein expression. Protein samples were combined with 2× Laemmli sample buffer containing 2-mercaptoethanol and heated at 75 °C for 10 min to denature proteins. Samples were then cooled on ice for 5 min and centrifuged at 13,280 rpm for 30 min to collect protein lysates. Proteins were separated by electrophoresis on a 10% SDS-polyacrylamide gel using Tris-glycine-SDS running buffer in a Bio-Rad mini-gel system. Electrophoresis was carried out at 80 V for 20 min followed by 155 V for 55 min. After separation, proteins were transferred onto nitrocellulose membranes using a Bio-Rad semi-dry transfer apparatus. Membranes were blocked with 5% milk prepared in PBS-T for 1 h at room temperature to reduce nonspecific binding. Membranes were then incubated overnight at 4 °C with the appropriate primary antibodies. Following primary antibody incubation, membranes were washed three times with TBS-T for 5 min each. HRP-conjugated anti-mouse or anti-rabbit secondary antibodies were then applied for 1 h at room temperature. Membranes were developed using Pierce ECL Western Blotting Substrate (Thermo Scientific). Protein bands were visualized and recorded using the iBright FL1000 chemiluminescence imaging system.

### Subcellular fractionation of mitochondria and cytoplasm

Subcellular fractionation was performed to isolate mitochondrial and cytoplasmic protein fractions. Cells were seeded onto 100 mm dishes and incubated for 24, 48, or 72 h post-infection. Following treatment, culture supernatant was collected, and cells were washed with cold PBS supplemented with Sigma 100× protease/phosphatase inhibitor cocktail diluted 1:50. Cells were lysed by adding 1 mL of mitochondrial homogenization buffer containing 10 mM Tris, 1 mM EDTA, and 250 mM sucrose, pH 7.4, supplemented with protease/phosphatase inhibitor cocktail. The buffer was evenly distributed across the dish to ensure complete coverage, and cells were incubated on ice for 10 min. The cell layer was then scraped and transferred into pre-chilled tubes. Samples were sonicated on ice using a probe sonicator with a 1.5 mm probe at 1% output for a total sonication time of 1.5 min using cycles of 3 s on and 30 s off for three cycles. Following sonication, samples were centrifuged at 10,000 × g for 10 min at 4 °C. A portion of the resulting supernatant was reserved as the whole-cell lysate fraction. After the whole-cell lysate fraction was collected, the pellet from this clarification step was discarded, and the remaining supernatant was transferred to pre-labeled tubes for mitochondrial isolation. The remaining supernatant was centrifuged at 12,000 × g for 20 min at 4 °C to pellet the mitochondrial fraction. The supernatant from this step was transferred to labeled tubes to represent the cytoplasmic fraction. The mitochondrial pellet was resuspended in the mitochondrial homogenization buffer supplemented with inhibitors and centrifuged again at 12,000 × g for 10 min to further purify the mitochondrial fraction. 2× Laemmli buffer containing β-mercaptoethanol was then added to the samples, which were subsequently loaded for western blot analysis with 20 µg of protein per lane. Fraction integrity was confirmed using organelle-specific markers, including tubulin for cytoplasmic and whole-cell lysate fractions and Tom70 for mitochondrial fractions. Anti-HCV Hep C cAg antibody was used to detect viral protein localization.

### Free fatty acid assay

Free fatty acid levels within each cell fraction were measured using the Free Fatty Acid Quantification Assay Kit (Abcam, ab65341) according to the manufacturer’s instructions. Briefly, lipids were extracted by mixing 25 µL of each fraction with 200 µL chloroform/Triton X-100 and incubating on ice for 30 min. Samples were centrifuged at 12,000 rpm for 5 min, and the lower organic phase was collected. The organic phase was air-dried at 50 °C in a fume hood to remove chloroform, leaving dried lipid residues. Lipids were then resuspended in 200 µL assay buffer. For fraction samples, 25 µL of resuspended lipid was mixed with 25 µL assay buffer, while 50 µL of supernatant samples were used directly. Positive controls were prepared using 6 µL fatty acid standard solution combined with 44 µL assay buffer, while negative controls consisted of 50 µL assay buffer. Following addition of 2 µL ACS reagent, samples were incubated at 37 °C for 30 min. Reaction mix was then added and samples were incubated for an additional 30 min at 37 °C protected from light. Absorbance was measured immediately at OD 570 nm using a microplate reader to quantify free fatty acid levels.

### Reverse transcription quantitative PCR

Reverse transcription quantitative PCR was performed to quantify gene expression during HCV infection. Human hepatoma-derived Huh7.0 cells were seeded in 6-well plates and cultured until approximately 70% confluent. Cells were infected with HCV JFH-1 and harvested at the indicated time points, with mock-infected cells serving as controls. Total RNA was extracted from infected and mock samples using the Direct-zol RNA Miniprep Kit (Zymo Research, R2052). RNA concentration and purity were determined using a NanoDrop spectrophotometer by measuring the A260/A280 ratio. Equal amounts of RNA were reverse transcribed into complementary DNA using the iScript cDNA Synthesis Kit (Bio-Rad, 1708891). Quantitative PCR was performed using SsoAdvanced Universal SYBR Green Supermix (Bio-Rad, 1725271) on a Bio-Rad CFX96 real-time PCR detection system. Each reaction contained 10–20 ng of cDNA, gene-specific primers, and nuclease-free water in a final reaction volume of 20 µL. Thermal cycling conditions consisted of initial denaturation at 95 °C for 3 min, followed by 40 cycles of denaturation at 95 °C for 10 s and annealing/extension at 60 °C for 30 s. Melt-curve analysis ranging from 65 °C to 95 °C was performed following amplification to verify amplicon specificity. Relative mRNA expression levels were calculated using the 2⁻ΔΔCt method. GAPDH was used as the endogenous control for normalization. Fold changes in transcript abundance were calculated relative to mock-infected samples. Data represent the mean ± SEM from at least three independent biological replicates.

### Seahorse XF Cell Mito Stress Test

Mitochondrial respiration was measured using the Agilent Seahorse XFp Analyzer and the Seahorse XF Cell Mito Stress Test assay. The Seahorse XF Cell Mito Stress Test was performed at 48 hpi according to the manufacturer’s protocol. Huh7.0 cells were seeded into Seahorse XFp cell culture miniplates at a density of 4,000 cells per well based on cell counter measurements. Cells were allowed to attach for 2 h, after which they were infected with HCV JFH-1 at a viral stock titer of 4.67 × 10⁵ PFU/mL and an MOI of 3, corresponding to 25.7 µL of viral inoculum per well. Mock-infected wells were included as controls. Cells were incubated for 48 hpi prior to metabolic analysis. The day before the assay, the Seahorse XFp sensor cartridge was hydrated according to the manufacturer’s instructions. Briefly, 400 µL of sterile water was added to each utility well, and 200 µL was added to each surrounding moat. The cartridge was incubated overnight in a 37 °C non-CO₂ incubator. On the day of the experiment, the hydration water was replaced with XF calibrant, and the sensor cartridge was equilibrated for 1 h in a 37 °C non-CO₂ incubator before the assay. Immediately prior to analysis, culture medium in each well was adjusted for Seahorse assay conditions. Of the initial 80 µL of growth medium present in each well, 60 µL was removed and replaced with 160 µL of XF DMEM assay medium, resulting in a final assay volume of 180 µL per well. Cells were incubated in a 37 °C non-CO₂ incubator to allow temperature and pH equilibration prior to measurement. Injection compounds were prepared fresh on the day of the experiment. Oligomycin was loaded into Port A to achieve a final well concentration of 1.5 µM. FCCP was loaded into Port B to achieve a final well concentration of 2.0 µM. Rotenone/antimycin A was loaded into Port C to achieve a final well concentration of 0.5 µM. Oxygen consumption rate measurements were recorded under basal conditions and following sequential injections of oligomycin, FCCP, and rotenone/antimycin A. These measurements were used to determine mitochondrial respiration parameters, including basal respiration, ATP-linked respiration, maximal respiration, spare respiratory capacity, proton leak, and non-mitochondrial respiration. Data were processed using Agilent Seahorse Wave software and used to compare mitochondrial respiration between mock-infected and HCV-infected cells.

### Reactive oxygen species (ROS) detection

Intracellular reactive oxygen species levels were measured using the ROS Detection Cell-Based Assay Kit using DCFDA (Cayman Chemical, 601520). Huh7.0 cells were seeded at a density of 5,000 cells per well and allowed to attach prior to infection. Cells were infected with HCV JFH-1 at a viral stock titer of 4.67 × 10⁵ PFU/mL at an MOI of 3, and samples were collected at 24, 48, and 72 hpi, with mock-infected cells serving as controls. At each time point, cells were incubated with DCFDA reagent, which is oxidized by intracellular ROS to produce a fluorescent signal proportional to ROS levels. Fluorescence intensity was measured using a microplate reader at the recommended excitation and emission wavelengths according to the manufacturer’s instructions. Relative ROS levels were calculated by comparing fluorescence values between mock- and HCV-infected samples at each time point.

### Antibodies

The following primary antibodies were used for immunofluorescence staining and western blot analysis: anti-HCV core antigen/Hep C cAg mouse antibody (Santa Cruz Biotechnology, sc-57800), anti-β-tubulin 12G10 antibody, anti-CD36 rabbit antibody (Abcam, ab252922), anti-DGAT1 antibody (ABIN7190465), anti-TIP47 (sc-390968), anti-LIPE/HSL rabbit antibody (Proteintech, 17333-1-AP), anti-LIPG antibody (Thermo Fisher Scientific, PA1-16799), anti-ATGL mouse antibody (Santa Cruz Biotechnology, sc-368278), anti-CPT1A antibody (ABIN7146833), anti-Tom70 antibody (Cell Signaling Technology, 65619S), anti-CIDEB rabbit polyclonal antibody (A17140), anti-perilipin-2 rabbit monoclonal antibody (A20843), anti-CIDE-C rabbit antibody (Proteintech, 12287-1-AP), and anti-CIDE-A rabbit antibody (Proteintech, 13170-1-AP). HRP-conjugated anti-rabbit IgG and anti-mouse IgG secondary antibodies were used for western blot analysis, and Alexa Fluor-conjugated secondary antibodies were used for immunofluorescence staining as indicated.

## STATISTICAL ANALYSIS

Statistical analyses were performed using GraphPad Prism 8.3.0 software (La Jolla, CA, USA). Data are presented as mean ± SEM unless otherwise indicated. Comparisons between two groups were performed using an unpaired two-tailed Student’s t-test. Comparisons among multiple groups were performed using one-way or two-way ANOVA followed by appropriate multiple-comparisons of tests, as indicated in the figure legends. Simple linear regression was used to assess relationships between continuous variables where indicated. Survival curves were compared using the log-rank test. A p value < 0.05 was considered statistically significant. Significance is indicated as follows: *p < 0.05, **p < 0.01, ***p < 0.001, ****p < 0.0001; ns, not significant. The definition of n is provided in each figure legend.

## Acknowledgements

This work was supported by grants from NIH/NIAID 1R16AI189452-01 (Q.T.), and the National Institute on Minority Health and Health Disparities of the National Institutes of Health (award number G12MD007597) (Q.T.). The funders had no role in the study design, data collection and analysis, decision to publish, or preparation of the manuscript.

